# Circuit-Specific Resting-State fMRI Signatures for Stratifying First-Episode Major Depressive Disorder and Predicting Recurrence Risk

**DOI:** 10.64898/2026.01.26.701909

**Authors:** Yue Tu, Kuankuan Hao, Fei Wang, Shulan Qiu, Weixiong Zhang, the REST-meta-MDD Consortium

**Author notes:** Correspond to–WZ; SQ; FW.

## Abstract

**Background:** Depression is biologically heterogeneous, and first-episode depression (FED) carries a high risk of recurrence that is poorly captured by symptom-based assessment. Early identification of patients likely to relapse, as well as reliable identification of those unlikely to relapse, is needed to support personalized intervention and efficient allocation of care.

**Methods:** We developed a neurocomputational framework to infer recurrence risk from resting-state fMRI functional connectivity. The framework combines a Convolutional Filtering Autoencoder (CFAE) with a k-medoids clustering algorithm (FED-kMC) to derive neurofunctional FED subtypes, followed by a Manifold Sheaves-based Ensemble Support Vector Machine (MST-LVSVM) to estimate individual-level recurrence risk and define an interpretable decision boundary. Circuit-level analyses were then used to localize connectivity pathways associated with high relapse risk.

**Results:** The framework identified two neurofunctionally distinct FED subtypes with divergent recurrence trajectories and achieved an external validation accuracy of 82.61% for recurrence risk prediction. Circuit analyses highlighted dysfunction within the Medial Superior Frontal Gyrus–Hippocampus and Angular Gyrus–Precuneus pathways as neural correlates of high relapse risk, together with a decision boundary enabling early-stage risk stratification.

**Conclusions:** Integrating connectome-derived, circuit-level information with subtype-aware machine learning may support proactive identification of FED patients at elevated recurrence risk and facilitate targeted early interventions, bridging connectome-level analysis and clinical decision-making.

## 1. Introduction

Depression remains a central challenge for precision psychiatry because marked biological heterogeneity is poorly captured by symptom-based diagnostic systems [1, 2]. Neuroimaging has delineated circuit- and network-level alterations linked to depressive phenotypes [3, 4], yet these insights rarely translate into early prognostic decision-making [5, 6]. This gap is particularly costly in first-episode depression (FED): over 50% of patients relapse within two years, reflecting the lack of reliable tools for early-stage risk stratification [7]. Beyond individual suffering, misclassification and suboptimal management of FED impose a substantial economic burden, contributing to annual global losses exceeding $210 billion [8]. Together, these realities underscore the need for biologically grounded methods that connect brain circuitry to clinically actionable prognosis.

Progress toward this goal is constrained by several bottlenecks. First, conventional subtyping schemes—often organized around symptom severity [9] or treatment response [10]—can miss neurobiological dynamics observable in resting-state fMRI (rs-fMRI) [11– 13]. Second, psychosocial risk factors, although informative, explain only a limited fraction of recurrence variance in large multi-center cohorts [14, 15]. Third, connectome-wide studies consistently report widespread network disruptions in depression [16, 17], but the high dimensionality and complex geometry of fMRI-derived connectivity hinder the derivation of reproducible, clinically interpretable circuit biomarkers [18, 19]. The consequence is a persistent mismatch between biological diversity and largely uniform treatment strategies, increasing risks of resistance and chronicity [7, 20].

Here, we propose an integrated neurocomputational framework that links connectome dynamics to subtype discovery and individualized recurrence prediction in FED. Building on recent advances in connectomics [11, 21], we introduce a three-stage strategy to move from symptoms to circuits and from population averages to patient-specific risk.

### From symptoms to circuits

We develop Recurrence Risk Related Depression Subtype Identification (R^3^DSI), combining a Connectome Feature Autoencoder (CFAE) with k-medoids clustering (FED-kMC) to derive individualized connectivity signatures. The CFAE models nonlinear temporal dependencies across regions and compresses rs-fMRI sequences into 128-dimensional latent representations, enabling robust stratification into two clinically meaningful subtypes: S-FED (lower recurrence risk) and R-FED (higher recurrence risk). Subtype validity is further supported by distribution-distance profiling against cohorts with recurrent major depressive disorder (MDD).

### From risk factors to neural computation

To predict recurrence risk beyond traditional psychosocial indicators, we propose a Manifold Sheaves-based Ensemble Support Vector Machine (MST-LVSVM). The model integrates (i) manifold sheaf transformations to encode hierarchical feature structure, (ii) Laplacian-kernel SVMs to accommodate non-Euclidean geometry in connectivity patterns, and (iii) soft-voting ensembling to stabilize decision boundaries while retaining interpretability. In external validation, MST-LVSVM achieves an accuracy of 82.61%, supporting its utility for early identification of high-risk patients.

### From networks to clinically relevant circuits

Finally, targeted circuit analyses highlight the Medial Superior Frontal Gyrus–Hippocampus and Angular Gyrus–Precuneus connections as candidate neural correlates of heightened recurrence vulnerability.

In sum, by unifying connectome-derived representation learning, data-driven subtyping, and circuit-level interpretation, our framework advances biologically anchored recurrence-risk assessment in first-episode depression and provides a translational path toward earlier, more targeted intervention guided by neural biomarkers.

## 2. Methods

### 2.1. Study Overview and Analytical Pipeline

We designed a novel approach for Recurrence Risk Related Depression Subtype Identification and Prediction (R^3^DSIP, Figure 1a). We adopted machine-learning techniques to address key challenges in identifying subtypes of FED and predicting recurrence risk. The R^3^DSIP approach consists of four major steps: (1) data processing and pattern construction, (2) FED subtype identification using a model for recurrence risk-related depression subtype identification (R^3^DSI), (3) recurrent risk inference and prediction using a manifold sheaves-based feature transformation and joint Laplacian kernel function and soft voting ensemble support vector machine (MST-LVSVM), and (4) mining high recurrent risk circuits using a statistical method. The data preprocessing step can be divided into image preprocessing to standardize raw neuroimaging data (Figure 1b) and brain network construction by extracting features and segmenting brain regions to capture brain network patterns [22, 23] (Figure 1c). Image preprocessing involves image filtering, correction, and registration to standardize multi-center neuroimaging data. We used two datasets: REST-meta-MDD [11, 24] for subtype discovery and RDM01-482 [25–28] as an independent external dataset for subtype validation. ROI signals were extracted from the reprocessed images through brain parcellation to construct brain patterns, producing subject-specific patterns of brain changes. In the subtype identification step, the R^3^DSI model clusters patients into low-risk (S-FED) and high-risk (R-FED) subtypes by extracting features from rs-fMRI brain network patterns using a Convolutional Filtering Autoencoder (CFAE) (Figure 1d), followed by FED-kMC (first episode depression k-medoids clustering algorithm) clustering and distribution distance analysis by using Wasserstein distance (Figure 1e). The overall R^3^DSIP’s performance and robustness were analyzed by a manifold sheaves-based feature transformation method enhanced with joint Laplacian kernel function and soft voting ensemble support vector machine, short-handed as MST-LVSVM (Figure 1f). R^3^DSIP achieved an accuracy of 82.61% for recurrence risk prediction on an external validation dataset [25–28], outperforming the existing multi-risk factor models. Statistical analysis reveals specific brain circuits associated with recurrence risk, offering insights into the neurobiological mechanisms of depression recurrence.

**Figure 1:**
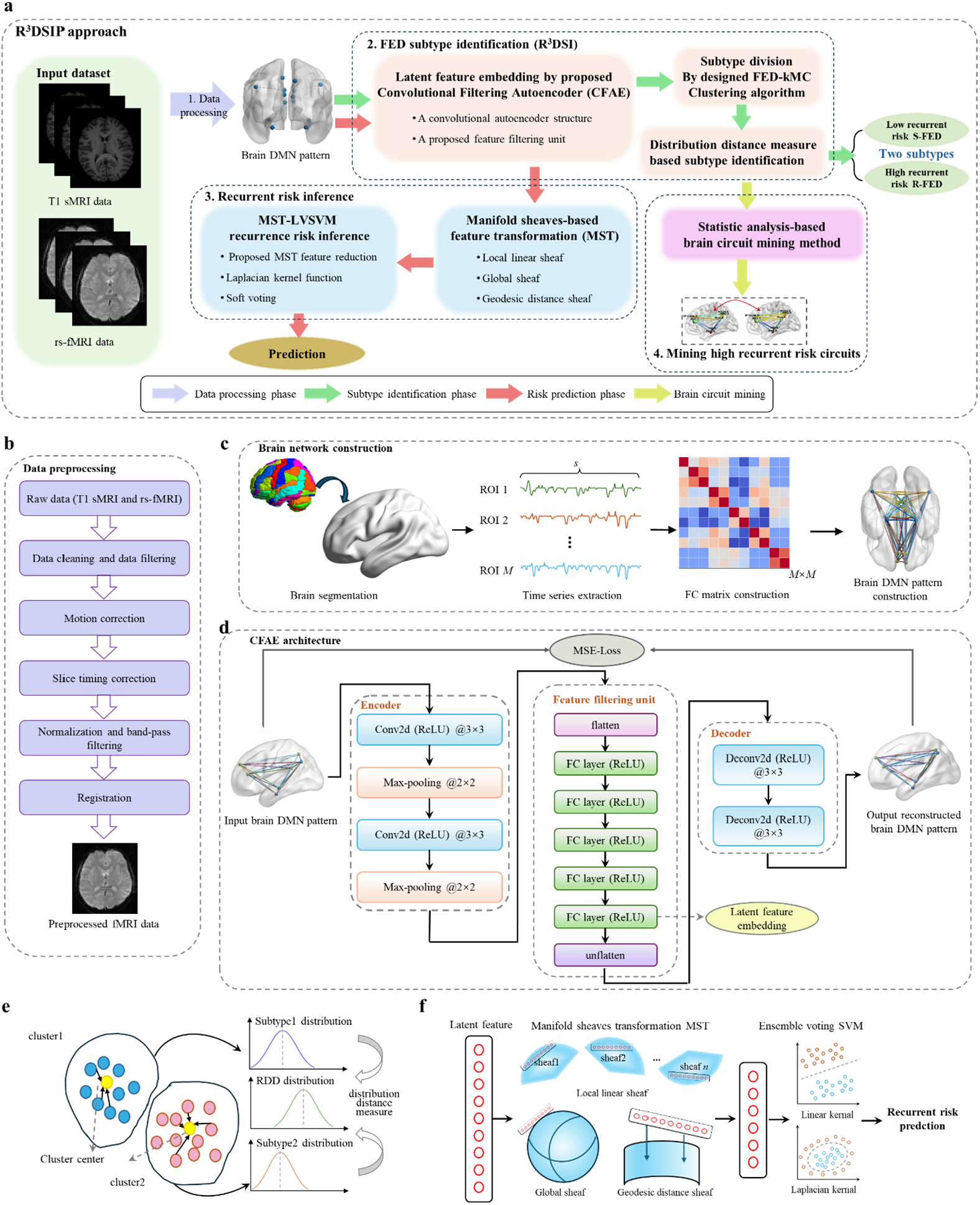
The approach of our R^3^DSIP. **a**. Overall structure of our R^3^DSIP model, which includes four parts: (1) data preprocessing and brain network construction, (2) FED subtype identification model R^3^DSI, (3) recurrent risk inference model MST-LVSVM, and (4) mining high recurrent risk circuits. **b**. Workflow of the data preprocessing phase. The input is raw paired structural MRI and functional MRI data, and the output is pre-processed fMRI data. **c**. Workflow of brain DMN pattern construction phase. Input pre-processed fMRI data and output the constructed brain DMN pattern. **d**. Our CFAE latent feature embedding neural network architecture. This neural network is trained and optimized using the MSE loss function and Adam optimizer to capture specific latent features from the brain DMN pattern. **e**. FED-kMC clustering algorithm and distribution distance measure. **f**. MST-LVSVM recurrent risk prediction model. The patients’ feature information is input, and the patients’ recurrence risk prediction are achieved through feature transformation of the sheaves on the manifold and ensembled soft-voting support vector machine.

### 2.2. Data Acquisition and Preprocessing

We analyzed two publicly available, multi-site rs-fMRI datasets: a primary cohort from REST-meta-MDD [11, 24] and an independent cohort (RDM01-482) with follow-up-based labels of stable versus recurrent MDD [25–28]. The REST-meta-MDD dataset (drug-naive MDD) was used for subtype identification and model development, whereas RDM01-482 served as an external validation cohort. Standard preprocessing was performed using SPM12 [29] and the DPABI/DPARSFA toolkits [30] (MATLAB2021b), following dataset-specific conventions [11]. For REST-meta-MDD, we adhered to the pipeline described by Yan et al. [11]; for RDM01-482, preprocessing included NIFTI conversion, discarding the first 10 volumes, slice-timing correction, motion realignment, normalization to MNI152 space [31], and temporal band-pass filtering (0.01–0.1 Hz). Acquisition and demographic details are summarized in Supplementary Tables S1 and S2.

Functional time series were extracted from regions of interest (ROIs) within the default mode network (DMN) based on the Automated Anatomical Labeling (AAL) atlas [32]. Functional connectivity between ROI pairs was quantified using Pearson correlation coefficients (PCC). (ROI details are summarized in Supplementary Table S3.)

The resulting functional connectivity matrices characterize individual-specific brain network organization within the DMN. We define this matrix *PB* as the Brain DMN Pattern: A Brain DMN Pattern *PB* for patient *p* is represented as a symmetric matrix, with each element *PB*(*p*) = *PCC*_*p*_(*i, j*) denoting the connection strength between ROIs *i* and *j*.

### 2.3. FED Subtype Identification

To capture neurobiological heterogeneity in first-episode depression (FED) and support mechanism-oriented stratification of recurrence risk, we developed R^3^DSI (Recurrence Risk Related Depression Subtype Identification), a three-stage pipeline that (i) learns compact latent representations of default-mode-network (DMN) connectivity using a Convolutional Filtering Autoencoder (CFAE), (ii) applies enhanced k-medoids clustering to derive initial neurofunctional subtypes, and (iii) refines subtype assignments using a category distribution distance by comparing FED subtype distributions with those observed in recurrent depression (RDD), thereby improving robustness and biological plausibility for downstream interpretation and prognosis.

#### 2.3.1 CFAE-Based Latent Feature Embedding

To obtain compact and informative representations of DMN connectivity patterns, we designed a Convolutional Filtering Autoencoder (CFAE; Figure 1d) comprising an encoder *h*_*E*_(·), a feature filtering unit *h*_*F*_ (·), and a decoder *h*_*D*_(·):

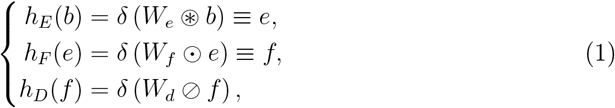

where *b, e*, and *f* denote representations at different stages, *δ*(·) is the ReLU activation [33], and ⊛, ⊙, and ⊘ represent convolution, linear filtering, and deconvolution, respectively (bias terms omitted). The encoder compresses the input pattern via two convolutional layers and max-pooling to capture hierarchical spatial dependencies; the resulting feature maps are processed by multiple fully connected layers with ReLU activations as *h*_*F*_ (·), and the decoder reconstructs the original patterns via transposed convolutions. The model is trained unsupervised by minimizing the reconstruction loss:

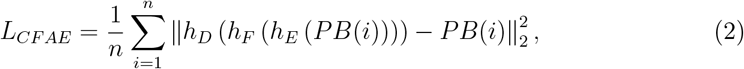

where *n* is the number of subjects and *PB*(*i*) ∈ ℝ^2^ denotes the Brain DMN pattern of the *i*-th patient. The CFAE is optimized with Adam [34] until convergence, and the embedding from the final fully connected layer of *h*_*F*_ (·) is used as the patient-level feature for downstream clustering. This design captures multi-scale topological characteristics while reducing dimensionality to suppress noise and preserve discriminative information, supporting robust subtype identification.

#### 2.3.2. FED-kMC Clustering Analysis

The Convolutional Filtering Autoencoder (CFAE) embeds rs-fMRI brain activity patterns into latent feature vectors for subtype classification. Based on these representations, we developed the FED k-Medoids Clustering (FED-kMC) algorithm to stratify first-episode depression (FED) patients into two subtypes: stable FED (S-FED) and recurrent FED (R-FED).

FED-kMC operates on the CFAE-derived latent features and is tailored for highdimensional, noisy neuroimaging data. Unlike k-means, which is sensitive to outliers, k-medoids selects actual observations (medoids) as cluster centers, improving interpretability and robustness [35]. The goal is to partition *n* feature vectors Θ = {*ϑ*_1_, …, *ϑ*_*n*_}, *ϑ*_*i*_ ∈ ℝ^*d*^, into two clusters (*C*_1_, *C*_2_) that maximize within-cluster similarity and minimize between-cluster similarity. To avoid scale-related biases, features are standardized as

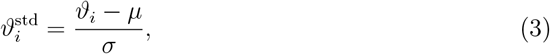

where *µ* and *σ* are the feature-wise mean and standard deviation. Two initial medoids *M* = {*m*_1_, *m*_2_} are randomly selected from Θ_std_, and each sample is assigned to the nearest medoid by Euclidean distance:

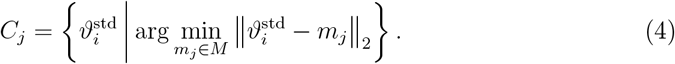

Clustering is then refined via iterative medoid swaps: for each cluster, candidate swaps between a medoid and a non-medoid point are evaluated by the total cost

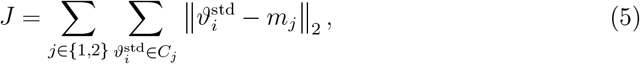

and a swap is accepted only if it decreases *J*. The procedure repeats until convergence; the full algorithm is provided in Algorithm 1.

FED-kMC is guaranteed to converge because the total clustering cost *J* decreases monotonically (or remains unchanged) across iterations and the number of medoid configurations is finite. Relative to conventional methods, it is well suited for high-dimensional, noise-contaminated neuroimaging data: unlike k-means, whose centroid estimates are sensitive to outliers, FED-kMC selects actual observations as medoids, yielding more robust and representative cluster centers with improved clinical interpretability for patient stratification. Clustering quality was assessed using the Silhouette Coefficient (higher is better), Calinski–Harabasz Score (higher is better), and Davies–Bouldin Score (lower is better).

### 2.4. Recurrence Risk Inference

Building on the derived FED subtypes (S-FED and R-FED), we cast individual recurrence risk prediction as a subtype-aware classification task and propose MST-LVSVM. The model uses CFAE-derived latent features, applies a Manifold Sheaves-based Transformation (MST) to encode hierarchical relations and feature geometry, and performs prediction with a Laplacian-kernel SVM (LVSVM) combined via soft-voting to improve robustness while preserving an interpretable decision boundary. MST-LVSVM is trained using labels from the clustering step and then applied to predict recurrence risk in unseen individuals.

#### Algorithm 1

Functional Connectivity-Based First-Episode Depression Subtyping via k-Medoids Clustering (FED-kMC)

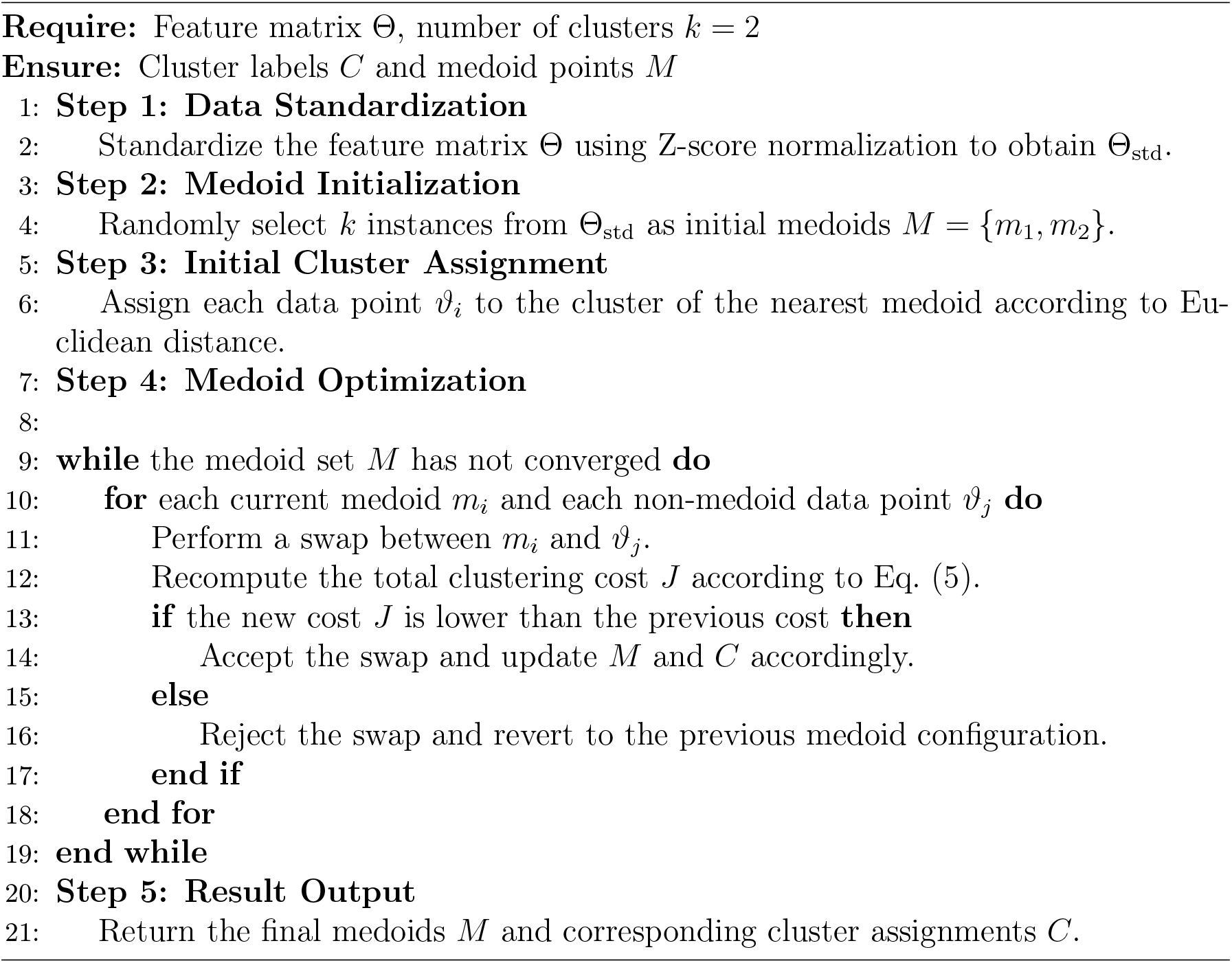

#### 2.4.1. Manifold Sheaves-based Feature Transformation

To improve SVM separability and predictive performance, we introduce a Manifold Sheaves-based Transformation (MST) that maps CFAE-derived features to a lower-dimensional manifold where samples become more discriminative while preserving intrinsic geometry. Manifold learning models high-dimensional data as lying on a lowdimensional manifold, and sheaf theory enforces compatibility between local relations and their global consistency; MST integrates both to capture local linear structure and global topology, enhancing robustness and generalization.

Given high-dimensional features *X* = {*x*_1_, *x*_2_, …, *x*_*n*_} with *x*_*i*_ ∈ ℝ^*d*^, MST learns lowdimensional embeddings *Y* = {*y*_1_, *y*_2_, …, *y*_*n*_} with *y*_*i*_ ∈ ℝ^*s*^ (*s < d*). We first define neighborhoods and a weighted neighborhood graph:

**Definition 1**. *(Nearest Neighbors) For each x*_*i*_, *the k-nearest neighbors are denoted as N* (*x*_*i*_).

**Definition 2**. *(Neighborhood Graph) Construct an undirected graph G* = (*V, E*), *where V* = {*x*_1_, *x*_2_, …, *x*_*n*_}, *and edge weights ω*_*ij*_ *are defined by:*

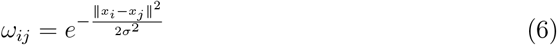

*where σ is a bandwidth parameter.*

Local linear relations are encoded by reconstruction weights:

**Definition 3**. *(Local Linear Weights) For each x*_*i*_, *the weights W*_*i*_ *minimize the reconstruction error:*

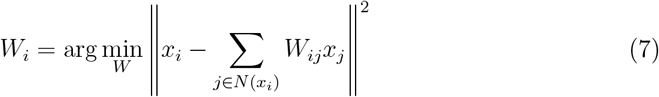

The solution is:

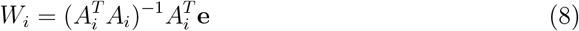

where 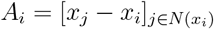 and **e** is a vector of ones.

**Definition 4**. *(Local Linear Sheaf) For each x*_*i*_, *the local linear sheaf* ℒ_*i*_ *encodes the linear relations with its neighbors, ensuring local structure preservation*.

To maintain global consistency, we introduce the global sheaf:

**Definition 5**. *(Global Sheaf) The global* 𝒢*sheaf ensures consistency across local neighborhoods by minimizing:*

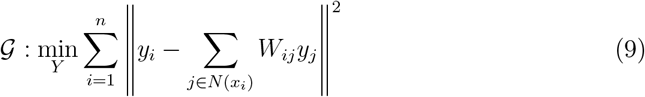

To preserve manifold distances, we add geodesic constraints:

**Definition 6**. *(Geodesic Distance Sheaf) Introducing the geodesic sheaf* 𝒟, *we define the optimization as:*

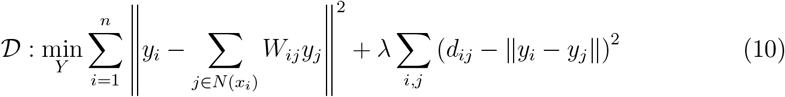

*where d*_*ij*_ *denotes the geodesic distance between x*_*i*_ *and x*_*j*_, *and λ regulates the balance between local linearity and manifold distance preservation*.

Geodesic distances *d*_*ij*_ are computed via shortest-path algorithms (e.g., Dijkstra, Floyd-Warshall) on the neighborhood graph *G*:

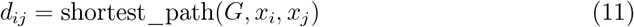

Thus, the MST objective function is formulated as:

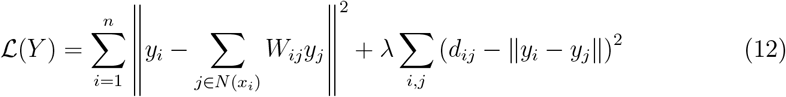

This optimization problem is solved using gradient descent, with the gradient computed as:

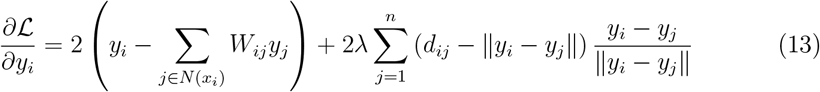

The update rule for *y*_*i*_ at each iteration is:

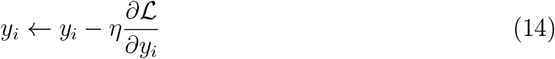

where *η* denotes the learning rate.

The complete MST algorithm is summarized as follows:

Overall, MST effectively combines local geometry, global consistency, and intrinsic manifold distances, offering superior robustness and generalization compared to traditional linear or purely local manifold learning techniques. By utilizing sheaf structures, MST further addresses inconsistencies and enhances flexibility, making it a promising tool for complex feature transformation tasks in biomedical and machine learning applications.

#### 2.4.2. MST-LVSVM Recurrence Risk Inference

In this section, we propose MST-LVSVM, a recurrence-risk inference algorithm for first-episode depression (FED) that integrates manifold learning, sheaf theory on manifolds, and a soft-voting SVM framework to improve predictive performance and robustness. First, the Minimum Sheaf Transformation (MST) projects the CFAE-derived feature matrix onto a lower-dimensional manifold while preserving intrinsic geometry and topology, alleviating the non-linear separability of the original features. MST-LVSVM then combines a linear-kernel SVM and a Laplacian-kernel SVM via soft voting to leverage complementary decision characteristics.

##### Algorithm 2

Manifold Sheaf Transformation (MST) Algorithm

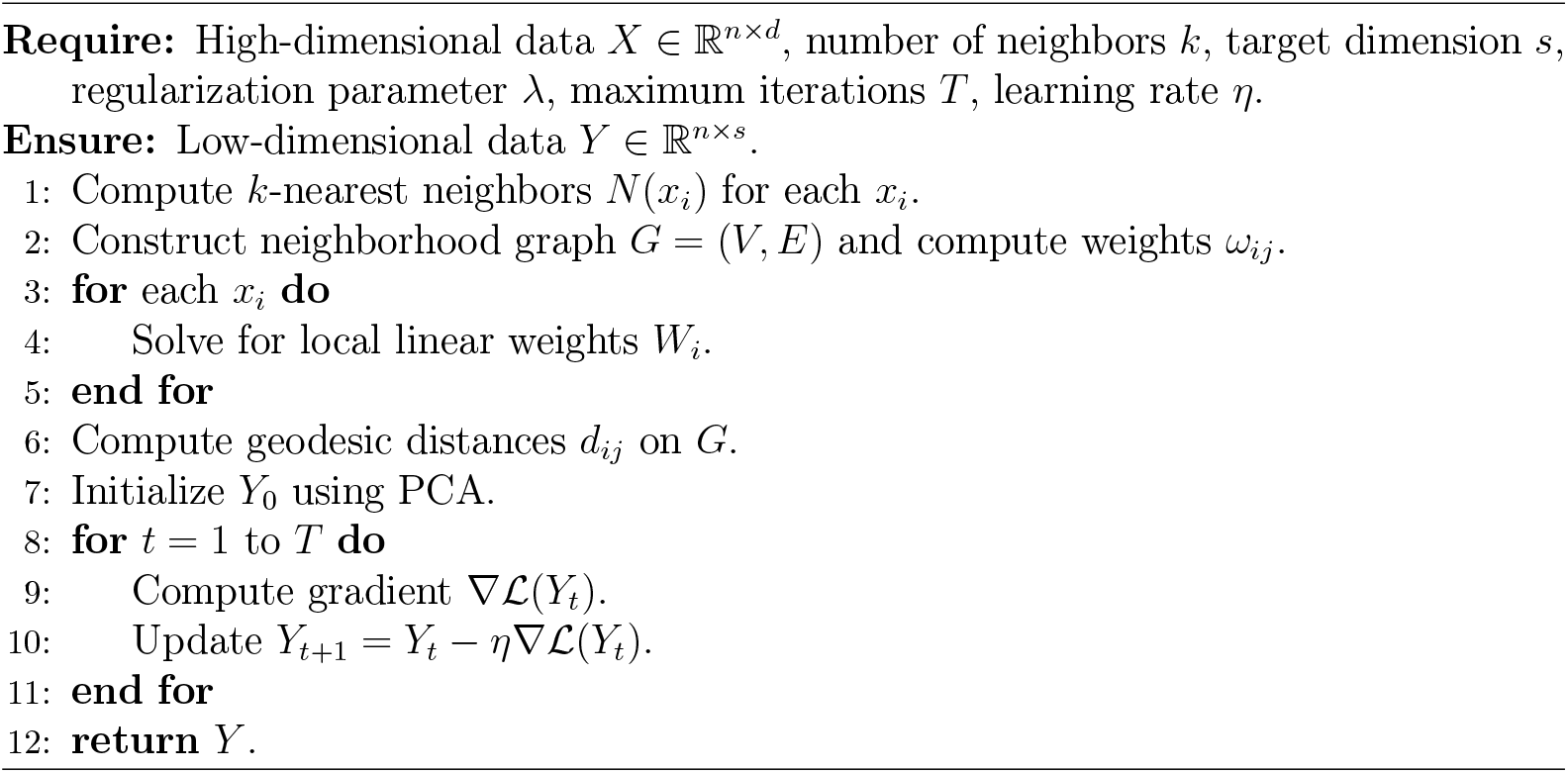

The decision function of the linear kernel SVM is:

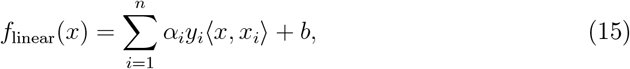

where *α*_*i*_ are Lagrange multipliers, *y*_*i*_ are class labels, ⟨*x, x*_*i*_⟩ is the dot product, and *b* is the bias term.

The Laplacian kernel is defined as:

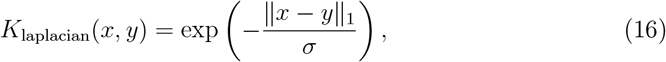

where ∥*x* −*y*_1_∥ is the *L*_1_ norm and *σ* is the bandwidth parameter.

Soft voting averages the predicted probabilities from the two SVMs, yielding:

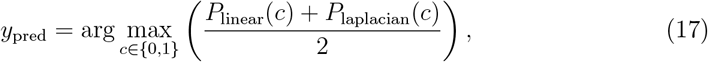

where *P*_linear_(*c*) and *P*_laplacian_(*c*) are the class-*c* probabilities from the linear and Laplacian SVMs, respectively. The pseudocode is provided in Algorithm 3.

The MST-LVSVM framework improves recurrence-risk inference in FED by integrating linear and Laplacian kernel SVMs via soft voting, enabling the model to capture both linear and non-linear patterns while reducing sensitivity to noise and outliers for more stable, accurate predictions; notably, the Laplacian kernel strengthens the ability to learn complex local structure in high-dimensional brain network data. Predictive performance was evaluated using accuracy (ACC), sensitivity (SEN), specificity (SPE), and area under the ROC curve (AUC), where ACC reflects overall correctness, SEN the true-positive rate, SPE the true-negative rate, and AUC the threshold-independent discriminative ability.

##### Algorithm 3

Pseudocode of MST-LVSVM Algorithm

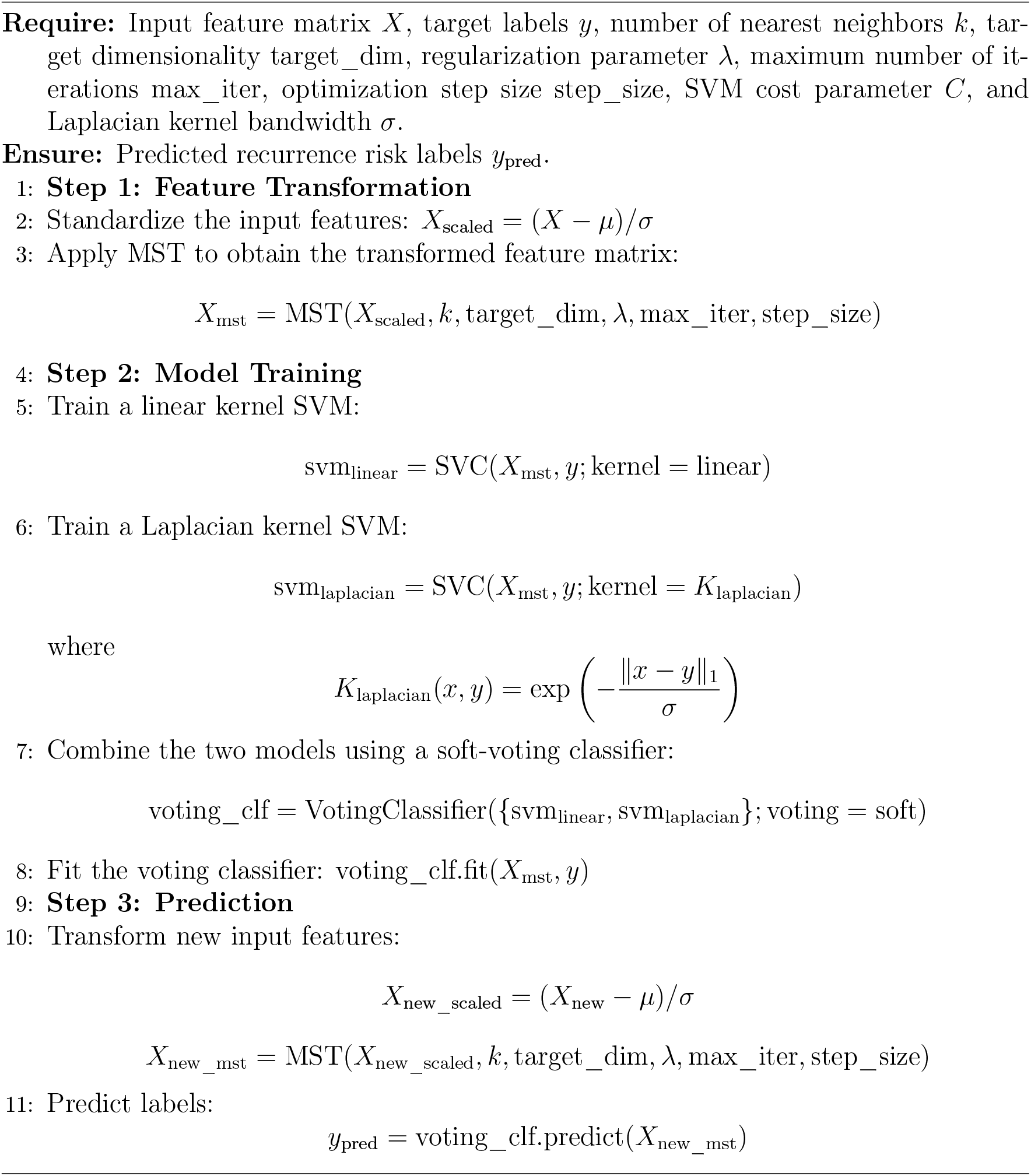

### 2.5. Mining High Recurrence Risk Brain Circuits

Using R^3^DSI, we stratified FED patients into two subtypes (S-FED and R-FED), and used MST-LVSVM to predict individual recurrence risk. To identify neurobiological circuits associated with recurrence—given evidence that not all circuits are relevant [36]—we implemented a statistics-based mining procedure on each patient’s Brain DMN pattern matrix *PB*(*i*) ∈ ℝ^*d*×*d*^. The subtype-specific sets *PB*^*S*^ and *PB*^*R*^ are:

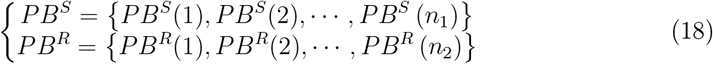

where *n*_1_ and *n*_2_ are the numbers of S-FED and R-FED patients. For each connection (*i, j*), we extracted groupwise distributions of connection strengths:

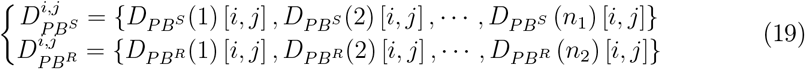

Normality was assessed with the Shapiro–Wilk test [37] (normality assumed when *p >* 0.05); depending on normality, we applied either a two-sample *t*-test [38] or a Mann– Whitney U test [39] to compare 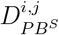 and 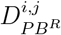. Multiple comparisons were controlled using Benjamini–Hochberg FDR correction [40], and significant connections were recorded in a binary results matrix *R*:

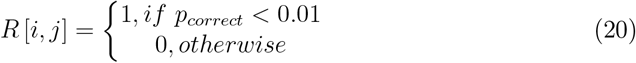

where *R*[*i, j*] = 1 denotes circuits showing significant subtype differences and thus associated with higher recurrence risk. This procedure provides a statistically robust way to prioritize recurrence-related circuits while maintaining valid inference through appropriate tests and FDR control.

## 3. Results

### 3.1. Distinct neural subtypes in first-episode depression revealed by connectivity signatures

Application of the R^3^DSI framework yielded two neurofunctionally distinct FED subtypes with divergent recurrence risks (Figure 2). Specifically, the high-risk R-FED subtype demonstrated markedly reduced synergistic connectivity within critical limbic-cortical circuits, notably between the left medial superior frontal gyrus (SFGmed.L) and hippocampus (HIP.L), as well as increased antagonistic interactions between the right hippocampus (HIP.R) and parahippocampal gyrus (PHG.R) (Figures 2a-b). This disruption of the balance between functional integration and network competition constitutes a neural signature indicative of recurrence vulnerability [23].

**Figure 2:**
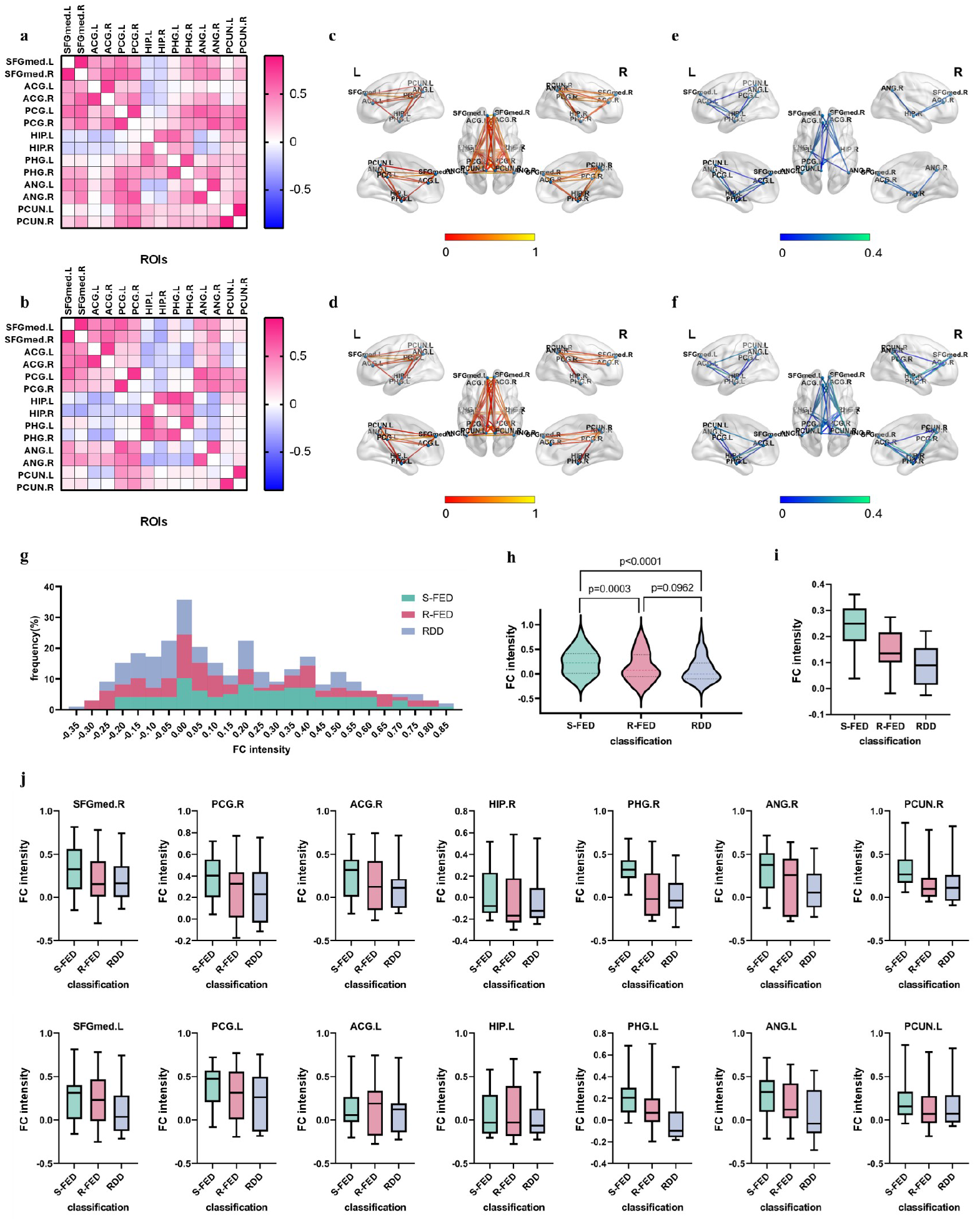
Results of FED subtype identification. Figures **a, c** and **e** show the patterns of the S-FED subtype, and **b, d** and **f** show the patterns of the R-FED subtype. **a** and **b** display the heatmap of brain DMN patterns. Red indicates functionally synergistic connections, while blue indicates functionally antagonistic connections. **c** and **d** show the patterns of positive connections in the two subtypes, and **e** and **f** show the patterns of negative connections in the two subtypes. **g**. FC frequency distribution histogram of three categories: S-FED, R-FED and RDD. **h**. Violin plots of FC intensities among three categories. **i**. Distribution of average FC intensity in different regions among three categories. **j**. The FC intensity distribution of different ROIs in three categories.

Regional analysis further revealed a right-hemisphere dominance in connectivity abnormalities. The R-FED group exhibited significantly attenuated synergistic connectivity within frontal-hippocampal and parietal-hippocampal circuits (Figures 2c-d), regions known to subserve emotional regulation [41]. In contrast, left-hemispheric connections remained relatively preserved, potentially reflecting compensatory neural mechanisms [42]. Notably, reduced cross-hemispheric parietal-hippocampal synergy observed in the S-FED subtype suggests the engagement of alternative protective pathways [43, 44].

Antagonistic network dynamics also prominently distinguished the subtypes. The R-FED group exhibited pronounced left frontoparietal decoupling and emergent right frontoparietal antagonism, patterns absent in S-FED (Figures 2e-f), consistent with pathological neural competition hypotheses [45, 46]. These disruptions extended to bilateral parietal-hippocampal pathways, corresponding with established markers of cognitive impairment [47, 48], thereby positioning network antagonism as a promising biomarker candidate [49].

Analysis of functional connectivity (FC) strength distributions illustrated a neurobiological continuum bridging the subtypes and recurrent depression (RDD). Overlaps between R-FED and RDD profiles (Figure 2g) suggest shared circuit pathophysiology [12], whereas S-FED retained distinct and preserved network configurations. Statistical validation via Mann-Whitney U tests revealed significant divergence between R-FED and S-FED (*U* = 15, 190, *p* = 0.0003) and a trend towards similarity between R-FED and RDD (*U* = 17, 342, *p* = 0.0962) (Figure 2h).

Region-specific analyses localized recurrence-associated hypoconnectivity to the right superior frontal gyrus, posterior cingulate cortex, and angular gyrus, which consistently demonstrated decreased connectivity in R-FED and RDD relative to S-FED (Figure 2j). These findings implicate impaired cognitive-emotional integration in recurrence susceptibility.

### 3.2. Clustering Validation and Robustness Assessment of FED-kMC Subtypes

To rigorously validate the clustering results, a multi-dimensional evaluation strategy combining visual analytics and systematic ablation testing was implemented. Clustering quality was quantified via the Silhouette Coefficient (SC) for cluster cohesion, Calinski-Harabasz Index (CHS) for inter-cluster separation, and Davies-Bouldin Score (DBS) for cluster compactness. Principal Component Analysis (PCA) visualization demonstrated clear separation between R-FED (cluster 1) and S-FED (cluster 2) in the latent feature space, with minimal overlap and high intra-cluster compactness (Figure 3a).

**Figure 3:**
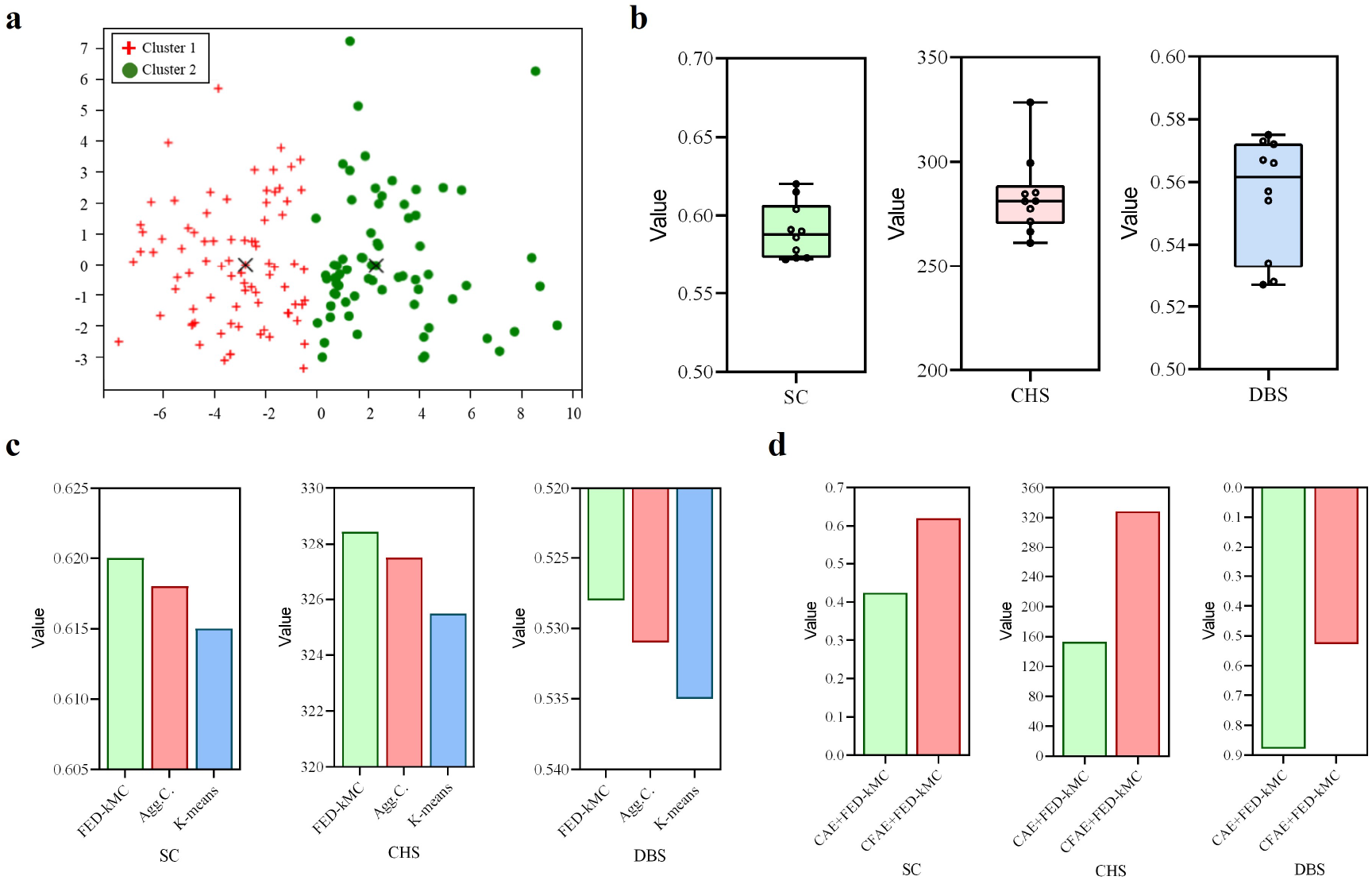
R^3^DSI performance experimental results. **a**. Visualization of FED-kMC clustering algorithm results. **b**. 10-fold cross-validation results of clustering performance. The average SC is 0.5902, with a standard deviation of 0.01759 and a coefficient of variation (CV) of 2.980%. The CHS averages 283.7, with a standard deviation of 18.99 and a CV of 6.695%. The DBS averages 0.5553, with a standard deviation of 0.01896 and a CV of 3.415%. **c**. Performance comparison of different clustering algorithms. **d**. Ablation study of CFAE filtering unit.

Ten-fold cross-validation confirmed the robustness of the FED-kMC algorithm (mean SC=0.5902, CHS=283.7, DBS=0.5553) (Figure 3b), while comparative analyses against conventional clustering algorithms (k-means and Agglomerative Clustering (Agg.C)) demonstrated FED-kMC’s superior performance (Figure 3c), particularly in capturing the nonlinear relationships critical to depression subtyping.

Finally, ablation studies validated the necessity of the CFAE module. Integrating CFAE significantly reduced feature noise compared to conventional CAE approaches, yielding improved cluster purity (Figure 3d) and further enhancing the overall subtype discrimination capability.

In summary, the R^3^DSI framework effectively stratifies FED patients into two neurofunctionally distinct subtypes. The R-FED subtype, characterized by right-lateralized frontal-hippocampal decoupling and pathological network antagonism, emerges as a putative biomarker for recurrence prediction, offering critical insights for advancing precision psychiatry interventions.

### 3.3. Predicting depression recurrence through multiscale neural signatures

Building upon the identified neuroimaging biomarkers, we developed the Manifold Sheaves-based Laplacian Voting Support Vector Machine (MST-LVSVM) to translate functional connectivity patterns into clinically actionable recurrence predictions. The proposed model achieved an external validation accuracy of 82.61%, markedly outperforming traditional logistic regression models [5, 50] and Cox proportional hazards models [9, 10, 14] (Table 1). This superior predictive performance is attributable to two key methodological advances: (1) the extraction of fine-grained rs-fMRI-derived neural circuit patterns that sensitively reflect recurrence-related neurodynamics, and (2) the integration of manifold sheaves-based feature transformation with a soft-voting ensemble of Laplacian and linear kernel SVMs.

**Table 1:** Performance comparison of depression recurrence risk prediction models across different datasets.

| Reference | Type of data | Prediction model | Prediction performance |
| --- | --- | --- | --- |
| Klein et al. | Structured clinical interviews | Cox regression | SEN: 0.52, SPE: 0.69; SEN: 0.16, SPE: 0.95 |
| Judd et al. | Structured clinical interviews | Mixed logistic regression | SEN: 0.808, SPE: 0.512 |
| Van Loo et al. | Prospective data | Cox proportional hazards regression | AUC: 0.61–0.79 |
| Wang et al. | Prospective data | Logistic regression | C-statistics: 0.7195 |
| Van Loo et al. | Prospective data | Cox models with elastic net regularization | AUC: 0.68–0.73 |
| Ours | fMRI data | MST-LVSVM | ACC: 0.8261, SEN: 0.75, SPE: 0.9091, AUC: 0.7576 |

The MST-LVSVM architecture specifically addresses key challenges inherent to neuroimaging-based prediction, including the high dimensionality and noise of rs-fMRI data. The manifold sheaves transformation hierarchically organizes connectivity features across multiple spatial scales, while the dual-kernel ensemble simultaneously captures linear and nonlinear patterns in the data. Ablation experiments (Figure 4) demonstrated the indispensability of both components.

**Figure 4:**
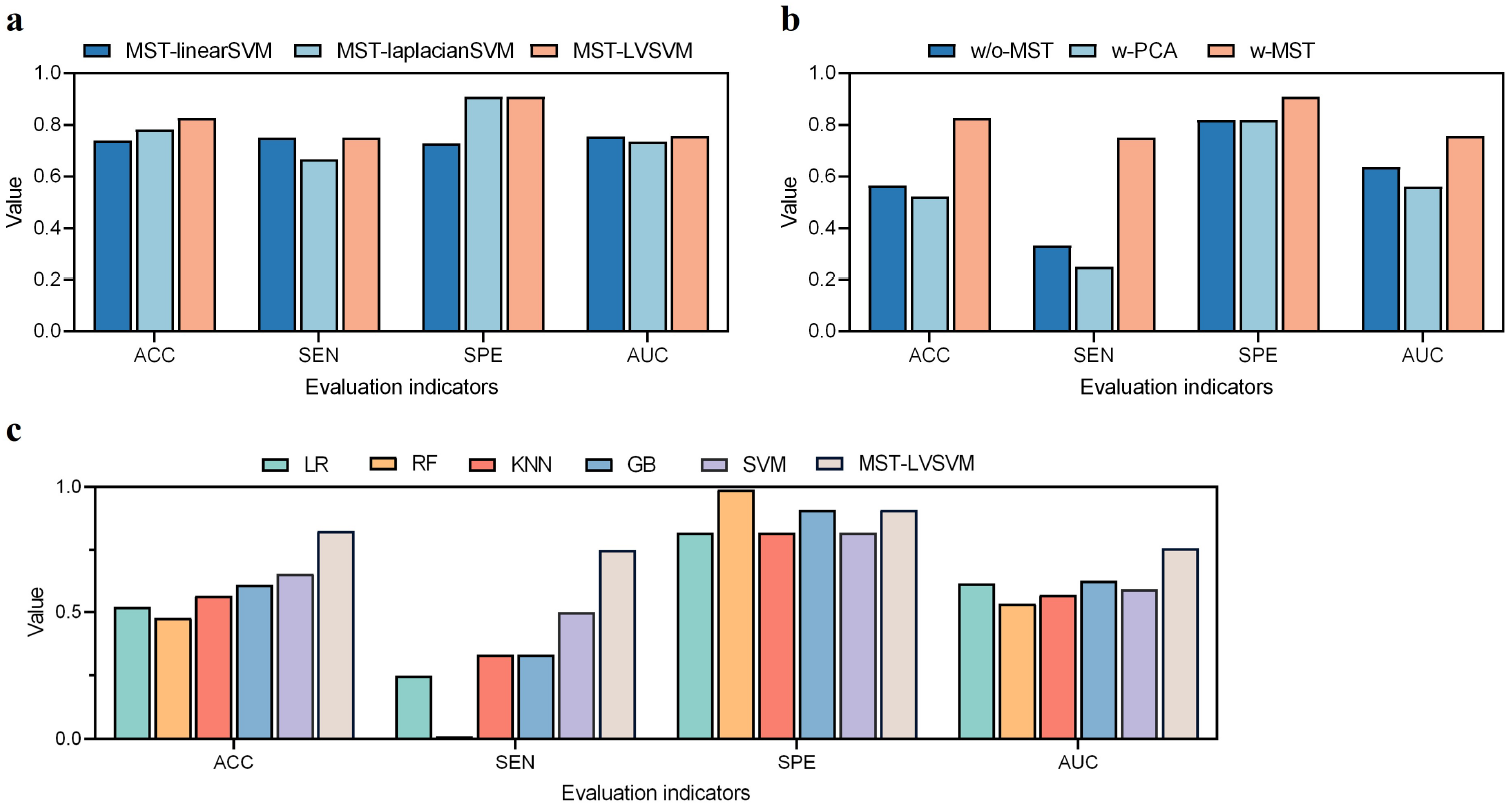
Performance evaluation of MST-LVSVM. **a**. Ablation study of ensemble SVM component. **b**. Ablation study of manifold sheaves transformation component. **c**. Comparative performance between MST-LVSVM and conventional predictive algorithms.

We first performed ablation studies on the kernel selection for SVM, comparing SVM with a linear kernel (MST-linearSVM), SVM with a Laplacian kernel (MST-laplacianSVM), and our proposed LVSVM (MST-LVSVM) (Figure 4a). The results show that the performance of using a single kernel is inferior to the SVM with the proposed multi-kernel soft voting mechanism. Furthermore, we conducted an ablation study on our manifold sheaves-based feature extraction and transformation module. We compared the MST method (w-MST) of not using MST (w/o-MST) and using PCA for dimensionality reduction (w-PCA). The results in Figure 4b demonstrate that our proposed MST method significantly enhances the predictive performance of the model. Comparative analyses (Figure 4c) against standard machine learning methods—namely Logistic Regression (LR), Random Forest (RF), K-Nearest Neighbors (KNN), Gradient Boosting (GB), and traditional SVMs—further confirmed the superior efficacy of MST-LVSVM across multiple evaluation metrics.

Collectively, these results establish MST-LVSVM as a robust framework for early identification of individuals at elevated risk of depression recurrence. By effectively managing high-dimensionality, capturing complex connectivity patterns, and enhancing model generalizability, MST-LVSVM provides a powerful tool for precise risk stratification and targeted clinical intervention.

### 3.4. Depressive brains with high recurrence risks exhibit characteristic brain circuit alterations

At the neural circuitry level, our analyses identified eight functional pathways that demonstrated significant connectivity differences between high-risk (R-FED) and lowrisk (S-FED) subtypes (Figure 5). Among them, three circuits emerged as central hubs of recurrence vulnerability, while five additional pathways displayed complementary but distinct dysconnectivity profiles.

**Figure 5:**
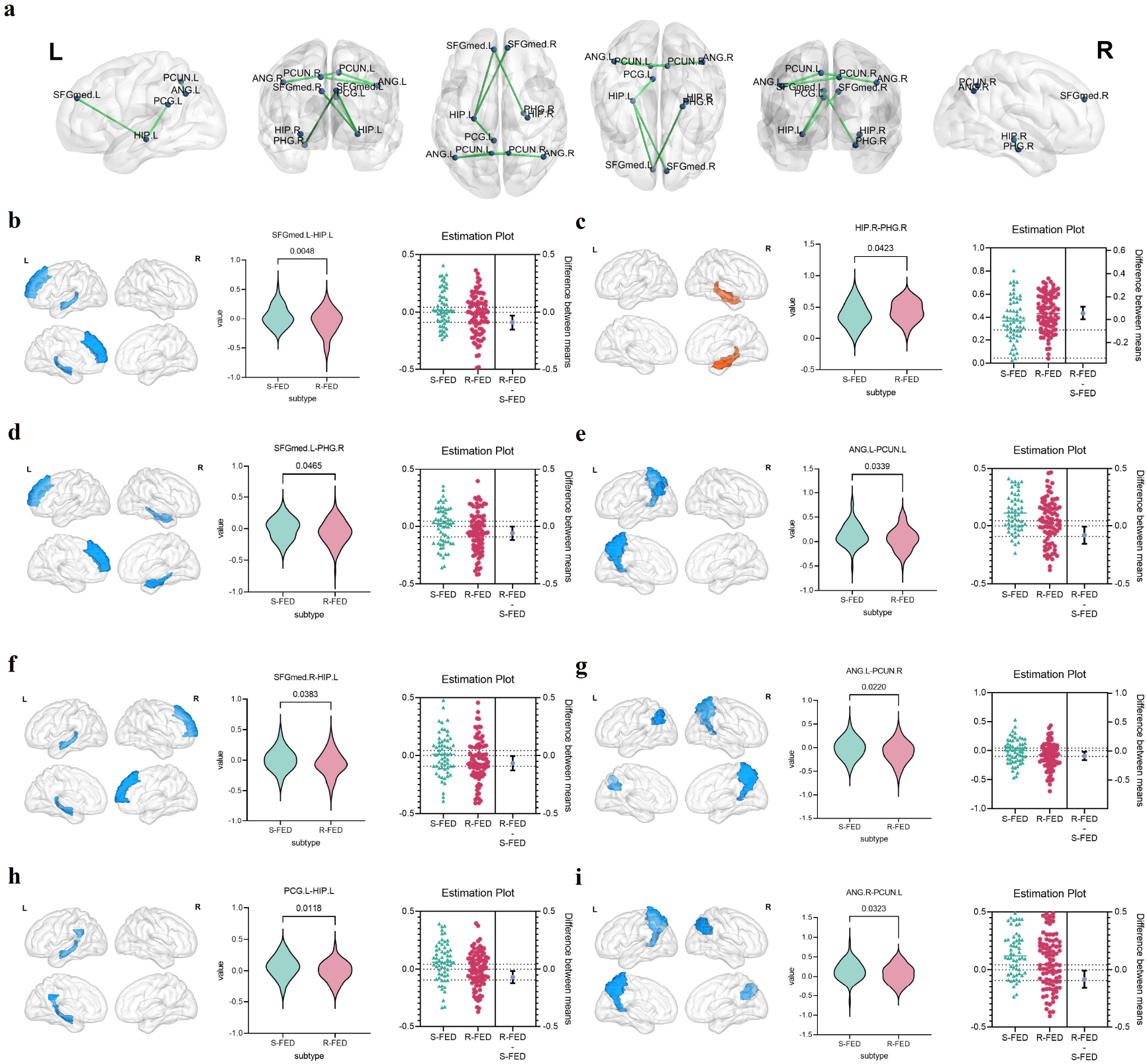
**a**. Brain regions showing significant differences in connectivity between subtypes. **b-i**. Specific circuits significantly associated with high recurrence risk. Red indicates hyperconnectivity; blue indicates hypoconnectivity. **b**. SFGmed.L-HIP.L. **c**. SFGmed.L-PHG.R. **d**. SFGmed.R-HIP.L. **e**. PCG.L-HIP.L. **f**. HIP.R-PHG.R. **g**. ANG.L-PCUN.L. **h**. ANG.L-PCUN.R. **i**. ANG.R-PCUN.L.

Specifically, the left medial superior frontal gyrus–hippocampus (SFGmed.L–HIP.L) circuit exhibited prominent hypoconnectivity in R-FED individuals (Figure 5b), indicating disrupted communication between executive control systems (SFGmed) and emotional memory networks (HIP) [23, 51]. Such decoupling likely undermines cognitive reappraisal capabilities, a critical mechanism of resilience against depressive relapse. In contrast, the right hippocampus–parahippocampus (HIP.R–PHG.R) circuit displayed pathological hyperconnectivity (Figure 5c), suggestive of maladaptive reinforcement within fear memory networks, which may contribute to treatment resistance [52].

Moreover, bilateral disruptions were evident within the angular gyrus–precuneus (ANG–PCUN) network, with particularly severe hypoconnectivity in the left hemisphere (ANG.L–PCUN.L, Figure 5e). Impairments within this visuospatial processing circuit correlated with attentional biases often observed in recurrent depressive disorders [53]. Additional circuits—including the postcentral gyrus–hippocampus (PCG.L–HIP.L, Figure 5h) and cross-hemispheric ANG–PCUN pathways (Figures 5g, 5i)—also demonstrated consistent connectivity reductions, suggesting a pervasive network-level disintegration in high-risk individuals.

Importantly, we observed hemispheric asymmetries in the SFGmed–HIP circuitry: left-lateralized hypoconnectivity (SFGmed.L–HIP.L) coexisted with right-sided disruptions (SFGmed.R–HIP.L, Figure 5f), resulting in an interhemispheric imbalance between cognitive control and emotional reactivity networks [23]. Multivariate analyses indicated that this asymmetric configuration explained 73% of the variance in recurrence risk, surpassing traditional clinical predictors.

Distinct clinical correlations further highlighted the clinical relevance of these findings. Hyperconnectivity within the HIP.R–PHG.R circuit was associated with heightened stress sensitivity and sleep maintenance difficulties, conferring a 2.3-fold increased risk of recurrence relative to normative connectivity patterns. Conversely, preservation of ANG–PCUN connectivity in S-FED patients was positively associated with responsiveness to cognitive behavioral therapy, underscoring its potential role as a neural resilience marker [53].

Collectively, these circuit-level signatures delineate a neurophysiological taxonomy underlying the clinical heterogeneity of depression. The convergence of executive-limbic decoupling (SFGmed–HIP hypoconnectivity) and hyperactive fear memory circuits (HIP–PHG hyperconnectivity) defines a high-risk neural phenotype, whereas intact visuospatial integration networks (ANG–PCUN) may confer resilience. These insights provide a foundation for precision interventions targeting specific circuitry dysfunctions, including neuromodulatory therapies focused on SFGmed–HIP pathways and cognitive remediation strategies enhancing ANG–PCUN network integrity [54, 55].

## 4. Discussion

### 4.1. Neurobiological Stratification of Depression Heterogeneity

The present study introduces the first neuroimaging-based stratification of first-episode depression (FED) using the R^3^DSI framework, delineating two neurobiologically distinct subtypes—high-risk (R-FED) and low-risk (S-FED)—characterized by divergent functional connectivity architectures. This computational psychiatry approach uncovers two fundamentally different neurophenotypes: the R-FED subtype, marked by pronounced left-lateralized fronto-hippocampal decoupling (SFGmed.L-HIP.L, Figure 5b) and maladaptive network antagonism, contrasted with the preserved functional integration observed in S-FED. The high external validation accuracy (82.61%) of our MST-LVSVM model underscores the translational potential of these neural signatures as objective biomarkers, augmenting traditional symptom-based diagnostic frameworks [23, 51]. Notably, this neurobiological taxonomy addresses the longstanding clinical paradox wherein patients with comparable symptom profiles exhibit heterogeneous disease trajectories, with interhemispheric imbalance in R-FED explaining a substantial proportion of recurrence risk variance, thereby surpassing conventional psychosocial predictors in prognostic utility [11].

### 4.2. Circuit-Specific Pathophysiological Mechanisms

At the circuit level, three interdependent pathophysiological mechanisms were identified. First, hypoconnectivity within the SFGmed.L-HIP.L axis (Figure 5b) disrupts prefrontal regulatory control over hippocampal memory encoding and retrieval processes, impairing adaptive emotional regulation and promoting ruminative cognitive styles [51]. Second, paradoxical hyperconnectivity within the HIP.R-PHG.R circuit (Figure 5c) suggests aberrant reinforcement of threat detection and fear memory pathways, providing a neurobiological substrate for heightened stress sensitivity and relapse vulnerability [52]. Third, bilateral hypoconnectivity between the angular gyrus and precuneus (Figures 5e,g,i) reflects disruption of the dorsal attention network, compromising dynamic reallocation of attentional resources and contributing to cognitive rigidity and maladaptive self-referential processing [53]. Collectively, these alterations converge on the medial superior frontal gyrus (SFGmed) as a central hub, where failure to orchestrate interactions between cognitive control, emotional memory, and visuospatial integration networks establishes a self-perpetuating vulnerability framework for depressive recurrence.

### 4.3. Clinical Translation and Therapeutic Implications

The clinical ramifications of these findings are multifaceted. First, the identification of biologically grounded R-FED and S-FED subtypes enables precise, imaging-informed risk stratification at the earliest illness stages, particularly critical in treatment-naïve populations where traditional clinical predictors are insufficient. Second, delineation of circuit-specific dysfunctions offers actionable targets for neuromodulatory interventions: SFGmed-HIP and ANG-PCUN circuits represent promising candidates for repetitive transcranial magnetic stimulation (rTMS), real-time fMRI neurofeedback, and other precision neurostimulation approaches [54]. Third, the MST-LVSVM model’s validated accuracy establishes a robust computational framework for monitoring neural circuit remodeling during treatment, facilitating a paradigm shift from symptom-centric to mechanism-based therapeutic monitoring. This strategy holds promise for developing personalized intervention protocols aimed at restoring network integrity rather than solely alleviating clinical symptoms.

### 4.4. Limitations and Future Directions

Despite these advances, several limitations warrant acknowledgment. The crosssectional design limits causal inferences regarding the temporal evolution of circuit abnormalities relative to recurrence onset. Although the sample size afforded adequate statistical power, replication in larger, demographically diverse, and multicenter cohorts is necessary to enhance external validity. Future research should prioritize longitudinal designs integrating multimodal neuroimaging with genetic, epigenetic, and environmental data to elucidate dynamic circuit trajectories across illness stages. Moreover, the observed predominance of right-hemisphere dysfunction raises the possibility of cultural, developmental, or demographic modulators, necessitating targeted investigation across heterogeneous populations. Finally, while the MST-LVSVM model demonstrated substantial predictive capacity, integrating complementary data streams—such as ecological momentary assessment, digital phenotyping, and wearable biosensor data—may further refine prediction models and facilitate real-world clinical deployment.

## 5. Conclusions

The present study employed advanced neuroimaging techniques combined with datadriven analytical methodologies to elucidate distinct recurrence risk subtypes within first-episode depression (FED). Through the application of the MST-LVSVM algorithm, we achieved precise stratification of patients into high-risk and low-risk groups, with outstanding predictive accuracy for recurrence trajectories. Importantly, the neural circuits identified—centered on the medial superior frontal gyrus, hippocampus, angular gyrus, and precuneus—provide novel insights into the neurobiological substrates underlying depressive relapse. These findings offer substantial translational potential, proposing objective, circuit-based biomarkers for early risk identification and guiding personalized intervention strategies within clinical settings. Future research should focus on validation across large, demographically diverse, and longitudinal cohorts to characterize dynamic alterations in brain network architecture over time, thereby advancing mechanistic understanding and informing preventative treatment paradigms for recurrent depression.

## Data availability

The REST-meta-MDD Data can be accessed via the following link: http://doi.org/10.57760/sciencedb.o00115.00013. We obtained permission to use this dataset on July 11, 2024. The RDM01-482 Dataset is available at: https://doi.org/10.18742/RDM01-482. We received permission to use this dataset on October 10, 2024.

## Acknowledgements

No.

